# Timing of readiness potentials reflect a decision-making process in the human brain

**DOI:** 10.1101/338806

**Authors:** Kitty K. Lui, Michael D. Nunez, Jessica M. Cassidy, Joachim Vandekerckhove, Steven C. Cramer, Ramesh Srinivasan

## Abstract

Decision-making in two-alternative forced choice tasks has several underlying components including stimulus encoding, perceptual categorization, response selection, and response execution. Sequential sampling models of decision-making are based on an evidence accumulation process to a decision boundary. Animal and human studies have focused on perceptual categorization and provide evidence linking brain signals in parietal cortex to the evidence accumulation process. In this exploratory study, we use a task where the dominant contribution to response time is response selection and model the response time data with the drift-diffusion model. EEG measurement during the task show that the Readiness Potential (RP) recorded over motor areas has timing consistent with the evidence accumulation process. The duration of the RP predicts decision-making time, the duration of evidence accumulation, suggesting that the RP partly reflects an evidence accumulation process for response selection in the motor system. Thus, evidence accumulation may be a neural implementation of decision-making processes in both perceptual and motor systems. The contributions of perceptual categorization and response selection to evidence accumulation processes in decision-making tasks can be potentially evaluated by examining the timing of perceptual and motor EEG signals.

## Introduction

Decision-making has been extensively studied using two-alternative forced choice (2AFC) tasks that incorporate multiple stages of information processing (Ratcliff et al. 2016). In these tasks, participants typically perceive a visual or auditory stimulus (perception), categorize the stimulus and select one of two responses (decision-making), and respond with a motor action (response execution). Behavioral data consisting of response time (RT) and accuracy have been modeled by a number of sequential-sampling models (SSM) including drift-diffusion models (DDM; Link and Heath 1975; Ratcliff and McKoon 2008), linear ballistic accumulator models (Brown and Heathcote 2008) and leaky competing accumulator models (Usher and McClelland 2001), among others. In these models, sensory information is first encoded, followed by an evidence accumulation process to a threshold that triggers motor planning and execution. When fitting DDMs to behavioral data, sensory encoding time and motor execution time are typically estimated using a single parameter labeled *non-decision time*. The remaining time, labeled here as *decision-making time*, is the period of evidence accumulation, which is treated as a unitary process in most sequential sampling models.

Both animal and human researchers have found neural markers of evidence accumulation in electrophysiology by focusing on perceptual processing in the parietal cortex (see Shadlen and Kiani 2013; Kelly and O’Connell 2013; O’Connell et al. 2018). For instance, direct recordings from parietal cortical neurons (found in the lateral intraparietal area; LIP) in monkeys have identified cells whose firing rates progressively increase prior to a decision and response (Roitman and Shadlen 2002; Huk and Shadlen 2005; Churchland et al. 2008). The changing firing rates of these neurons are consistent with the theoretical account of accumulation of evidence to a boundary that is central to decision-making in SSMs. Moreover, in these studies, as the strength of sensory signals increases, the rate of increase in firing rate is enhanced, suggesting a faster rate of information accumulation that leads to faster RTs and more accurate decisions (Roitman and Shadlen 2002).

These findings in animal models have motivated studies in humans using EEG to identify a signal that ramps to a decision threshold, providing clear information about the rate and timing of decision-making (Kelly and O’Connell 2013; Philiastides et al. 2014). A number of studies have focused on the P300, a stimulus-locked evoked potential that increases its positive amplitude over parietal electrodes, reaching a maximum *at least* 300 milliseconds (ms) after stimulus presentation (Philiastides et al. 2006; Ratcliff et al. 2009) or the closely related central-parietal positivity (CPP), a label that better accounts for the variability in the timing of the peak of positive potentials across different experiments (O’Connell et al. 2012; Kelly and O’Connell 2013; Rangelov and Mattingley 2020). The P300 and CPP amplitudes are sensitive to stimulus probability and stimulus salience (Polich et al. 1996; Smith and Ratcliff 2004), such that low-probability and high-salience sensory events elicit higher amplitude signals. Moreover, the parietal signals have been observed for both auditory and visual stimuli, suggesting that they are supramodal responses (Polich et al. 1996; O’Connell et al. 2012) related to perceptual categorization (Duncan-Johnson and Donchin 1977; Kutas et al. 1977; Kotchoubey and Lang 2001; Azizian et al. 2006).

Both the amplitude and the timing of the CPP and P300 have been suggested as indicators of the evidence accumulation process. In a study using stimuli with different levels of salience and a categorical discrimination (face/car), the magnitude of the P300 parietal signal was correlated to the evidence accumulation rate in a DDM of the response data (Philiastides et al. 2006). In another study using vigilance tasks, the detection of gradual reductions in stimulus contrast evoked a CPP that was delayed as RT increased and peaked at the time of response execution (O’Connell et al. 2012). Critically, this signal was found across sensory modalities, specifically auditory and visual (Polich et al. 1996; O’Connell et al. 2012) and even in decision-making tasks not requiring a motor response (O’Connell et al. 2012). A similar effect could be observed for motion discrimination, with a delayed and smaller CPP peak amplitude as motion coherence decreased (Kelly and O’Connell 2013). This relationship has also been found for more complex perceptual tasks such as mental rotation (van Ravenzwaaij et al. 2017).

Decision-making tasks that require a motor response (such as a button press) have at least two decision-making components: *perceptual categorization* and *response selection*. Sequential sampling models such as the DDM divide the interval between stimulus and response into three phases: perceptual encoding, evidence accumulation, and response execution (Link and Heath 1975; Ratcliff and McKoon 2008). DDM models of behavioral data are limited to a single non-decision time parameter that lumps together perceptual encoding and response execution. Similarly, the DDM models perceptual categorization and response selection by a single evidence accumulation process that crosses an evidence threshold, triggering response execution. Previous studies described above have focused primarily on perceptual categorization and identified the P300/CPP as a correlate of evidence accumulation. Other studies who have focused on response selection by examining the readiness potential (RP), which is recorded over motor areas and has been interpreted as activity related to response selection in preparation for, and sometimes been found to be distinct from, response execution (Eimer 1998; Miller et al. 1999; Leuthold et al. 2004; van Boxtel and Böcker. 2004; Alexander et al. 2016). Previous studies have found that the RP can be elicited during evidence accumulation. Gluth et al. (2013) employed a paradigm that allowed a decision to be made about taking a stock at any point throughout six trials. They found that the stimulus-locked lateralized RP (LRP) was elicited in trials when participants did not take the stock option. Multiple other studies have found LRPs before stimulus onset, when pre-cued information is given about the response to be made, reflective of response selection (Osman et al. 1995; Leuthold et al. 1996; Ulrich et al. 1998; Leuthold et al. 2004). A response-locked RP can be observed before motor execution and is also influenced by pre-cued information, with the RP peak latency to response completion shorter when cued information about the finger to be used is provided (Rohrbaugh and Gaillard 1983; Osman et al. 1995). These studies suggest that the RP can be reflective of multiple movement-related processes, potentially including *response selection*.

We propose that depending on task demands, both perceptual categorization and response selection can contribute to the time course of decision-making. We carry out a new experiment that focuses on response selection and identify the RP as a motor signal that reflects the time course of evidence accumulation in a DDM model of RT. We sought to test the idea that response selection can reflect the time course of evidence accumulation. In this exploratory study, we make use of the DDM to model behavioral data in a 2AFC task known as the Action Selection (AS) task (O’Shea et al. 2007). In the AS task, the participants are presented one of four different stimuli varying along two dimensions (shape and size) and respond with one of two simple motor responses (shoulder rotations). The stimulus to response mapping in this task is thought to be more difficult than typical 2AFC tasks because it requires integration of two features, shape and size, in order to select the correct response. We make use of a DDM to separate the additive contributions of *non-decision time* (NDT: for perceptual preprocessing and motor execution) and *decision-making time* (DT) to *response time* (RT) while participants perform the AS task. The DDM makes use of a unitary evidence accumulation process to model DT, which we hypothesize incorporates both perceptual categorization and response selection. We demonstrate that in this particular task the RP, rather than the P300/CPP, will track DT. This close relationship suggests that the RP reflects an evidence accumulation process for response selection and supports the notion that evidence accumulation is a general neural implementation of decision-making in both perceptual and motor systems. Moreover, concurrent EEG recordings can be used to refine our understanding of the contribution of perceptual categorization and response selection to decision-making time.

## Methods

### Participants

Fifteen adults participated in this study. All participants met the following inclusion criteria: at least 18 years of age, right-handed, and English-speaking. Right-hand dominance was verified using the Edinburgh Handedness Inventory (Oldfield 1971). Participants were not able to participate if they demonstrated any of the following exclusionary criteria: inability to maintain attention or understand verbal instructions; any major neurological, psychiatric, or medical disease; or a coexisting diagnosis impacting arm/hand function. One participant was removed from analysis due to the EEG being contaminated pervasive muscle artifact, which precluded estimation of slow-wave potentials (i.e. the RP and P300), resulting in a sample size of fourteen adults (age 18-26 years; 9 female). This study was approved by the University of California, Irvine Institutional Review Board. Each subject provided written informed consent.

### Procedure

Participants performed two different tasks that required evaluation of a visual stimulus and a motor response. Each sat with their back on a chair, hips and knees at approximately 90 degrees, and right forearm in a tabletop splint. Participants completed 4 blocks (40 trials per block) of the AS task and 4 blocks (40 trials per block) of the Execution Only (EO) task (see below). The blocks alternated for all participants, starting with AS task. After each block, participants received a 30 second rest break that was prompted by a black screen on the laptop (Fig. 1b).

**Fig. 1:**
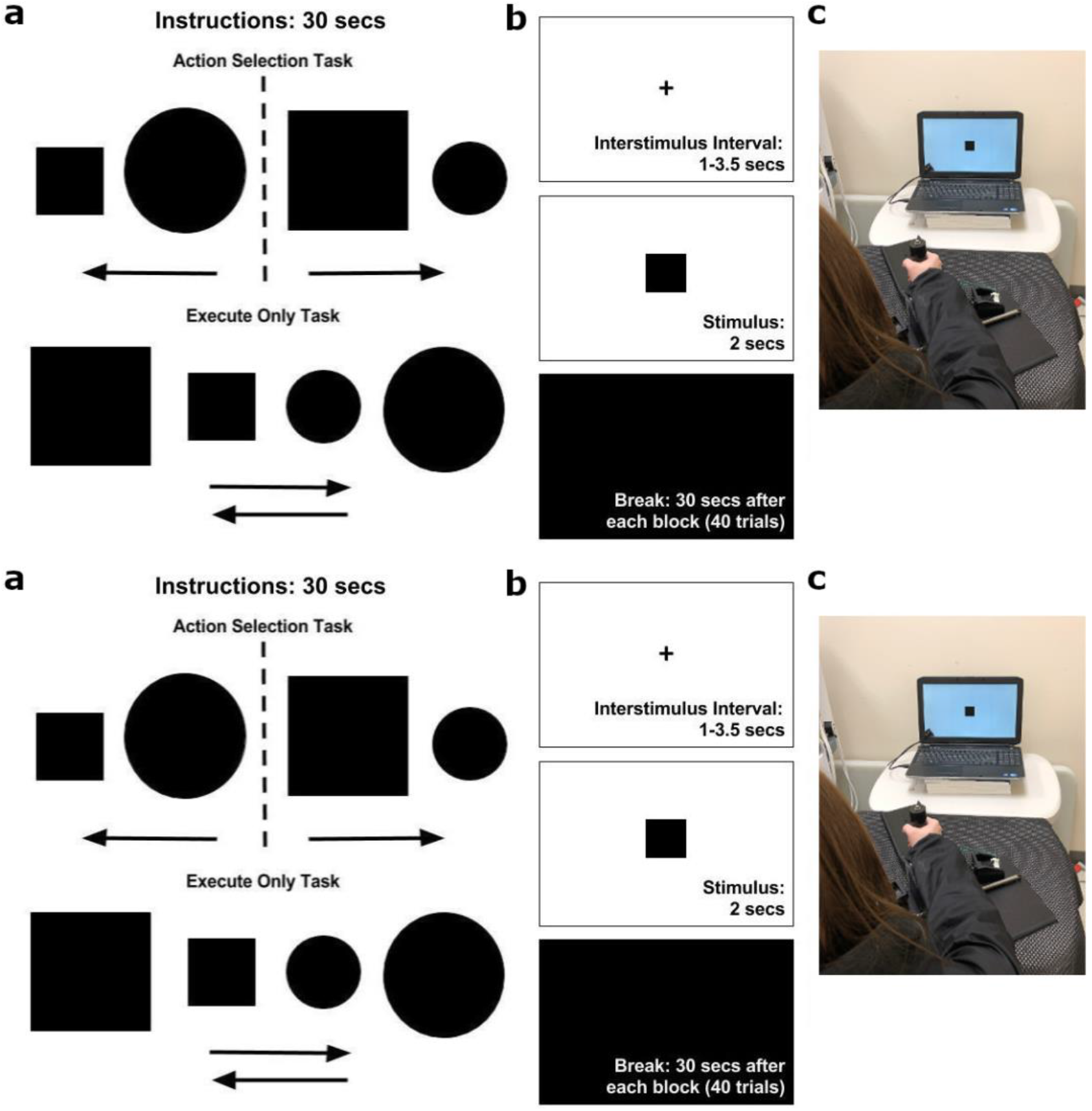
a) Participants received instructions prior to each block of 40 trials. The Action Selection (AS) task instructions directed the participants to make an external rotation movement when a large square or small circle appeared, and an internal rotation movement when a large circle or small square appeared. For each Execution Only (EO) block, instructions directed participants to perform either an external rotation movement or an internal rotation irrespective of the stimulus. b) The time course of each trial. The participants fixated on a cross during an interstimulus interval of random duration ranging from 1 to 3.5 seconds. A single stimulus was presented for two seconds during which responses were collected. After each block of 40 trials, participants received a 30 second break. c) The participants used the lower arm splint apparatus to make an internal (left) or external (right) shoulder rotation of 17.5 degrees to press a switch that captured their response. The splint was used to minimize any forearm or hand movements.

Each block of trials began with an instruction presented for 30 seconds (Fig. 1a). On each trial (Fig. 1b), participants focused on a fixation cross. A stimulus (i.e., shape) appeared for two seconds and the response was given by internal (inward) or external (outward) rotation of the right shoulder by 17.5 degrees using the tabletop forearm splint apparatus (Fig. 1c), until the splint made contact with a button embedded in either lateral wall. The purpose of the splint apparatus was to minimize compensatory movements by forearm and/or hand and thereby limit movement to a single direction in a single joint (i.e., right shoulder rotation). We used this splint apparatus in order to replicate this study design in stroke patients, where participants would likely be limited in making fine motor movements with distal limbs such as the fingers. This device enables gross motor movements through internal and external shoulder rotations. Participants could respond as soon as the stimulus was presented, and up to 3 seconds after stimulus presentation. Participants’ responses were recorded when they depressed the button, indicating movement completion. The interstimulus interval was randomized between 1 to 3.5 seconds. Participants received verbal instruction to move their arm to the middle of the apparatus after each trial.

### Experimental Tasks

The AS task (O’Shea et al. 2007) is a 2AFC task. Participants were instructed to perform an external right shoulder rotation (move the splint outward) when they saw either a big square or small circle and instructed to perform an internal right shoulder rotation (move the splint inward) when they saw either a big circle or small square. The large shapes spanned 2 degrees of visual angle while the small shapes spanned 1 degree of visual angle.

The EO task is a simple reaction time task that was used as a control condition, which generated all of the behavioral data and evoked potentials without any *necessary* perceptual categorization or response selection. Participants were instructed to move their right forearm, via shoulder rotation, to only one side within a given block, upon stimulus onset, irrespective of the size and shape presented. The EO blocks alternated between performing only internal rotations and performing only external rotations.

### Drift-Diffusion Model of Response Time

The RT and choice data for all participants and both tasks (AS and EO) were simultaneously fit to a hierarchical drift-diffusion model (DDM), using a Bayesian estimation method with Markov Chain Monte Carlo samplers (Plummer 2003; Vandekerckhove et al. 2011; Lee and Wagenmakers 2013; Waberisch and Vandekerckhove 2014). Fitting parameters of DDMs adds to the analysis of human behavior by assuming simple underlying cognitive processes that have some empirical validation (Voss et al. 2004). DDMs also add to the cognitive interpretation of EEG and fMRI signals by relating cognitive parameters to observed cortical dynamics (Mulder et al. 2014; Nunez et al. 2015; Turner et al. 2015; Nunez et al. 2017; Turner et al. 2017; Nunez et al. 2019a).

In a DDM of decision-making, it is assumed that humans accumulate evidence for one choice over another in a random-walk evidence-accumulation process with an infinitesimal time step until sufficient evidence is accumulated to exceed the threshold (labeled *boundary separation*) for either the correct or the incorrect choice. That is, evidence accumulates following a Wiener process (i.e. Brownian motion) with an average rate of evidence accumulation (labeled *drift rate*) until enough evidence for a correct decision over an incorrect decision is made (see Ratcliff et al. 2016 for a further discussion). The instantaneous variance (labeled *diffusion coefficient*) of this Wiener process was fixed at 1.0 in this study, because only two of the three evidence dimension parameters can be uniquely estimated (Ratcliff 1978; Ross 2014; Nunez et al. 2017). That is, the drift rate and boundary separation can always be scaled by the diffusion coefficient and produce the same fit of data.

Our hierarchical model was fit such that we could estimate separate model parameters for each participant derived from their accuracy-RT data. Each cognitive parameter of each participant was drawn from a single task-level population (AS or EO). This meant that we assumed that drift-diffusion cognitive parameters (drift rate *δ*, boundary separation *α*, and non-decision time *τ*) varied across participants (*p*) and the two tasks (*t*), but that there was similarity across the participants within each task. Adding these hierarchical parameters (mean parameters *µ* and standard deviation parameters *σ*) to summarize participants in each task yields better estimates of parameters due to shrinkage, a phenomenon whereby parameters are better estimated because hierarchical relationships enforce similarity across each participant (Gelman et al. 2014). Task-level Bayesian priors for hierarchical drift-diffusion model parameters were wide normal distributions. The prior distributions were centered at 1 for drift rate and boundary separation with standard deviations of 2 and 0.5 respectively. These two parameters are scaled by arbitrary evidence units dependent upon the scale imposed by the choice of fixed diffusion coefficient (here fixed at 1). The prior distribution for non-decision time τ was centered at 300 ms with a standard deviation of 250 ms and truncated at 0 ms. Note that we did not include start point of evidence accumulation parameters (see Ratcliff et al. 2016) because we found parameter estimates based on accuracy-RT and not left/right-RT data, and thus assumed that the start point of evidence accumulation was the midpoint between a correct and incorrect choice.

To refine our estimates of model parameters, we modeled the presence of contaminant trials to remove them from the decision-making model. Contaminant trials are defined as RT observations that are not due to an evidence accumulation process and are due to another random process. Because these RTs are not due to a decision process, the associated accuracy on those trials should be about 50% in a 2AFC task. DDM parameters are not well estimated in the presence of contaminant trials, so like previous implementations (Drugowitsch et al. 2012; Nunez et al. 2019a) we assumed that every participant in each condition (AS or EO) had some proportion (*λ*) of trials that were generated with a random response (Bernoulli distribution with probability parameter equal to 0.5) and a random (Uniformly distributed) RT between 0 and a large RT (bounded by the maximum observed RT for that participant in that particular task). The overall hierarchical model of accuracy and RT data (vector ***y***) per trial (*n*) is defined by the following normal (*N*), truncated normal (with truncation between *a* and *b* denoted by ∈ (*a, b*), gamma (*Γ*), and uniform (*U*) hierarchical, prior, and likelihood distributions:

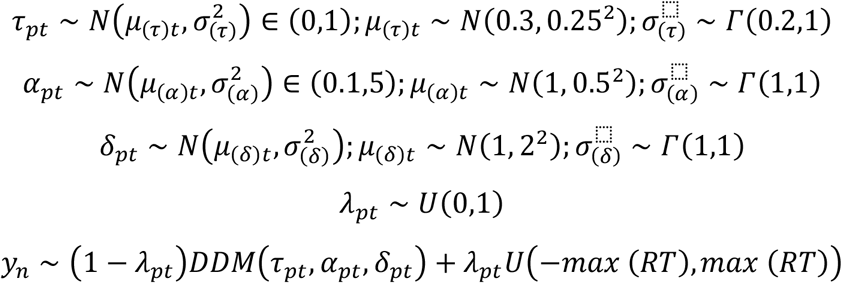

*RT* is defined as the amount of time needed after the stimulus onset for the participant to depress one of the two lateral wall buttons with the forearm splint apparatus. *DT* is the amount of time it takes to accumulate evidence to threshold (modeled by the random walk process), while *NDT* is the amount of time not associated with evidence accumulation, such as *visual encoding time* (i.e. the amount of time that the brain takes to recognize that evidence must be accumulated after visual onset; see Nunez et al. 2019a) and *motor execution time* (i.e. any time after decision-making occurs but before the response is recorded by the computer, including arm movement time in the apparatus).

For this study, the primary analysis focused on the posterior distributions of NDT for the two tasks instead of all estimated parameters from the hierarchical DDM because our hypotheses involved only the DT and NDT per participant and not the underlying shape of the DT and NDT distributions. Medians of posterior distributions of NDT were used as parameter estimates and were subtracted from RTs on each trial to obtain estimates of DT for each trial. Parameter estimates and 95% credible intervals (given by the 2.5th and 97.5^th^ posterior percentiles) of the hierarchical DDM are given in Table 2 of the Supplementary materials. The hierarchical DDM’s ability to describe data is provided in Table 1 of the Supplementary materials as summarized by in-sample prediction of correct-RT percentiles in each task.

### EEG Recording

The participants were fit with a 256-electrode EEG cap (HydroCel Sensor Net, Electrical Geodesics, Inc., Eugene, Oregon, USA). The cap was placed on the participant’s head after 10 practice trials of the AS task were completed to familiarize them with the experimental procedures. EEG data were sampled at 1000 Hz using a high-input impedance Net Amp 300 amplifier (Electrical Geodesics, Inc.) and NetStation 4.5.3 software. The EEG signals were referenced to the vertex electrode (Cz) during recording. The inputs from the splint apparatus were recorded by the EEG amplifier, using separate channels to record respective buttons for internal and external shoulder rotation movements. The onset of the stimulus was recorded by the EEG amplifier using a light detector (Cedrus, San Pedro, California, USA) for precise timing information synchronized with the EEG.

### EEG Preprocessing

All EEG pre-processing and analysis was performed using original MATLAB (Natick, MA) programs. The EEG data were detrended and segmented into trials starting 1000 milliseconds (ms) prior to stimulus onset up to 2500 ms after the stimulus for a total duration of 3500 ms. Any trials where the participants either responded incorrectly (wrong rotation direction), did not respond at all, or responded either too rapidly (within 200 ms) or too slowly (more than 2.3 seconds) from the stimulus onset, were not included in analyses. Out of the total 4,480 trials, for all participants in both tasks, there was only one trial that was considered too fast and six trials that were too slow. If participants responded more than once in one trial, the first response was considered their response. Bidirectional autoregressive interpolation was performed on each EEG channel from 5 to 30 samples after the response on each trial to remove an artifact created in the EEG by the input signal from the switches on the splint apparatus. As all data analysis took place in the interval prior to the response, this interpolation did not affect any of the results and was only performed to facilitate visualizing the results. The EEG data was high-pass filtered at 0.25 Hz (0.25 Hz pass, 0.1 Hz stop, 10 dB loss) and notch filtered at 60 Hz (pass below 59 Hz and above 61 Hz, 10 dB loss) using Butterworth filters.

The EEG data were artifact edited in two stages to remove some stereotypical artifacts generated from events such as eye movement, jaw movements, or muscle activity (Nunez et al. 2016). In the first stage, by visual inspection, trials that contained any vigorous movement and channels frequently containing EMG artifacts were removed from further data analysis. After the manual inspection, the data were re-referenced to the common-average reference. In the second stage, Independent Component Analysis (ICA) was then used to classify and remove artifacts from data that were related to eye movements, electrode pops, and environmental noise. ICA components were assessed as clearly artifact or possibly containing EEG (Nunez et al. 2016). All components not marked as artifact were kept for data analysis, and only components that clearly captured artifacts such as eye blinks, eye movements, or isolated channel pops were removed. The ICA components were then projected back into channels and the data were low-pass filtered at 50 Hz (Butterworth filter, 50 Hz pass, 60 Hz stop, 10 dB loss).

### Evoked potentials

For each participant, *stimulus-locked* evoked potentials (EPs) were calculated at each channel by aligning each trial to the marker of the stimulus onset and averaging across trials, and *response-locked* EPs were calculated by aligning to the marker of the button press and averaging across trials. In preliminary analyses, only a trivial difference was found in RT between internal and external shoulder rotations (see Results). Thus, we combined trials for the two directions of rotation to analyze the behavioral data and compute the EPs.

The stimulus-locked EPs that were analyzed were the N200, P300, and stimulus-locked RPs. In order to isolate each EP component, without contributions from other EPs, the P300 and stimulus-locked RPs were low-pass filtered at 4 Hz (Butterworth filter, 4 Hz pass, 8 Hz stop, 10 dB loss; Leuthold et al. 1996; Miller et al. 1999). In these EPs, the results are presented from 400 ms before the stimulus to 1500 ms after the stimulus, and baseline correction was performed by subtracting the mean potential 200 to 0 ms before the stimulus onset. The N200 waveforms were low-pass filtered at 10 Hz (Butterworth filter, 10 Hz pass, 20 Hz stop, 10 dB loss) and high-pass filtered at 1 Hz (Butterworth filter, 1 Hz pass, .25 Hz stop, 10 dB loss; Nunez et al. 2019a). These results are presented from 400 ms before the stimulus to 1500 ms after the stimulus, and baseline correction was performed by subtracting the mean potential 100 ms to 0 ms before the stimulus onset.

The only response-locked EP analyzed is the response-locked RP, which was low-pass filtered at 4 Hz (Butterworth filter, 4 Hz pass, 8 Hz stop, 10 dB loss; Leuthold et al. 1996; Miller et al. 1999). For the response-locked RPs, the results are presented from 1200 ms before the response to 100 ms after the response, with baseline correction performed by subtracting the mean potential in the interval 1200 to 1000 ms before the response.

### Evoked Potentials with Response Time Tertile Split

EPs were also calculated based on each participant’s RTs divided into three equal-sized bins to separate them into different response speed conditions (i.e., fastest, middle, slowest) for both tasks. The stimulus-locked EPs (P300 and stimulus-locked RPs) were calculated the same way as mentioned above. The response-locked RPs in the fastest and middle conditions also followed the same methods as mentioned above. However, in order to accurately estimate the onset times for the response-locked RPs in the slowest condition, the time window had to be extended to 1500 ms before the response to account for RTs that were longer than 1200 ms. Baseline correction was then performed by subtracting the mean potential in the interval 1500 to 1300 ms before the response. This time interval could not have been applied to the middle and fastest RT conditions because the response-locked RP would have potentially overlapped with data from the previous trial due to short interstimulus intervals (as short as 1000 ms, see Procedure above).

### Timing of the Readiness Potential: RP duration and RP onset time

We characterized the timing of the RP in terms of duration of the response-locked RPs and the onset time of the stimulus-locked RPs. The response-locked RP was calculated by averaging the EEG data aligned to the response. We identified the eight channels (out of 256) that displayed the strongest negative potential prior to the motor response, which is the defining characteristic of the RP. These eight channels were located close to the midline and slightly left-lateralized over motor areas of the brain (see Fig. 3c). RP duration, which was calculated from the response-locked RP, was defined as the interval during which the potential remained consistently negative at these channels, starting from the initial negative deflection up to the response completion. To identify this interval, the first derivative of the RP was estimated by taking the difference of the potential at each time point from the previous time point. The start of the response-locked RP was detected at the center of the first interval where the derivative remained negative for 125 ms, indicating a consistent negative deflection of the EP. The duration of the RP was defined as the interval from this starting point to the button press. The onset time of the RP relative to stimulus presentation was calculated from the stimulus-locked RPs at the same eight channels with the same requirement that the derivative must have remained negative for 125 ms to locate the onset of the negative deflection. Note that we also explored calculating response-locked and stimulus-locked RPs with 100 ms windows for negative derivatives. These changes did not fundamentally change the analysis results. However, the 100 ms windows did not capture obvious changes in the potentials that could be seen visually, so we chose to use the 125 ms window for both response-locked and stimulus-locked RPs.

**Fig. 2:**
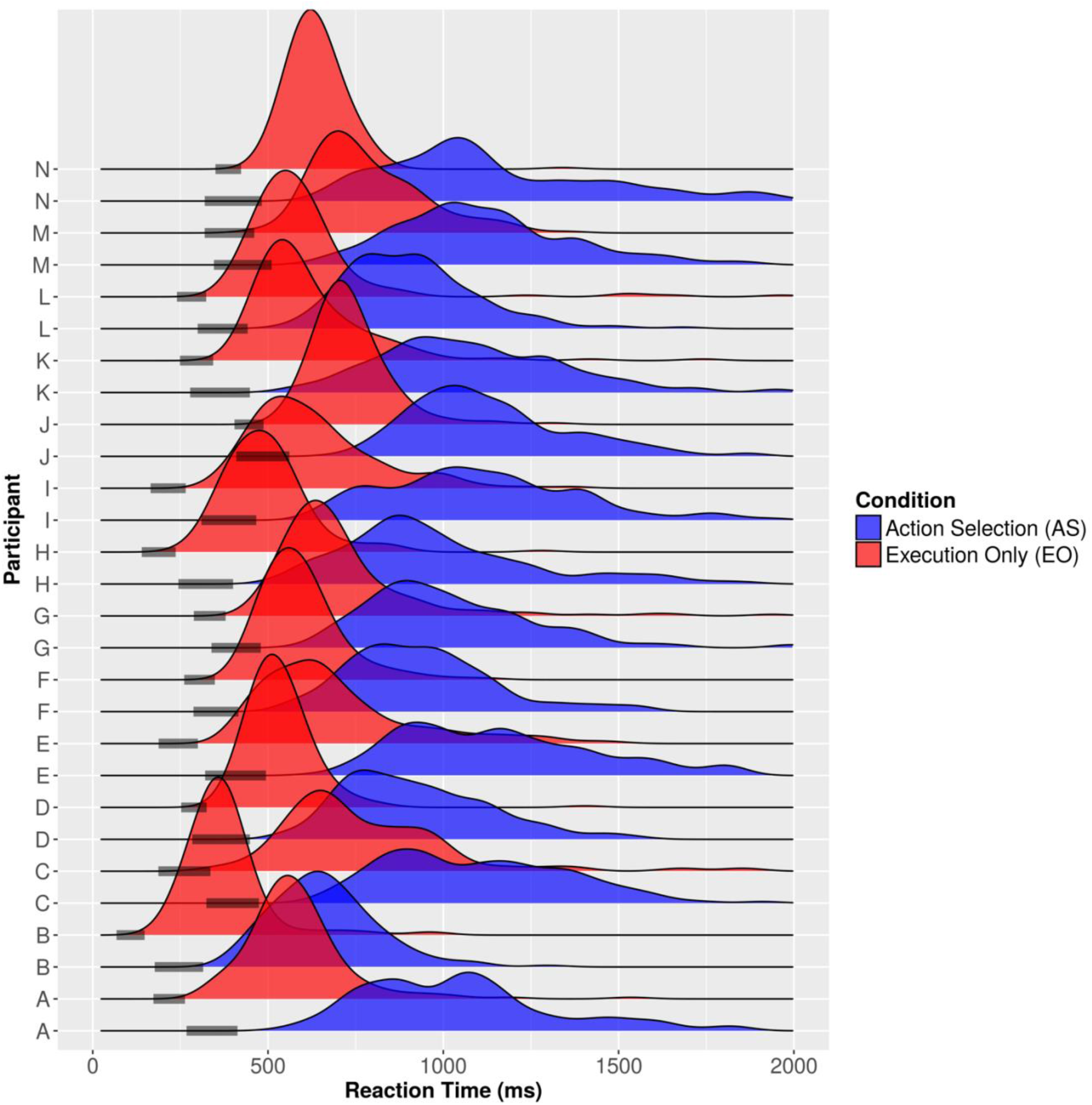
Response time distributions and estimated non-decision times for each participant. Each participant had a distribution of response times for both the AS and EO tasks. 95% credible intervals of non-decision time are given by the shaded bars on the response time axis.

**Fig. 3:**
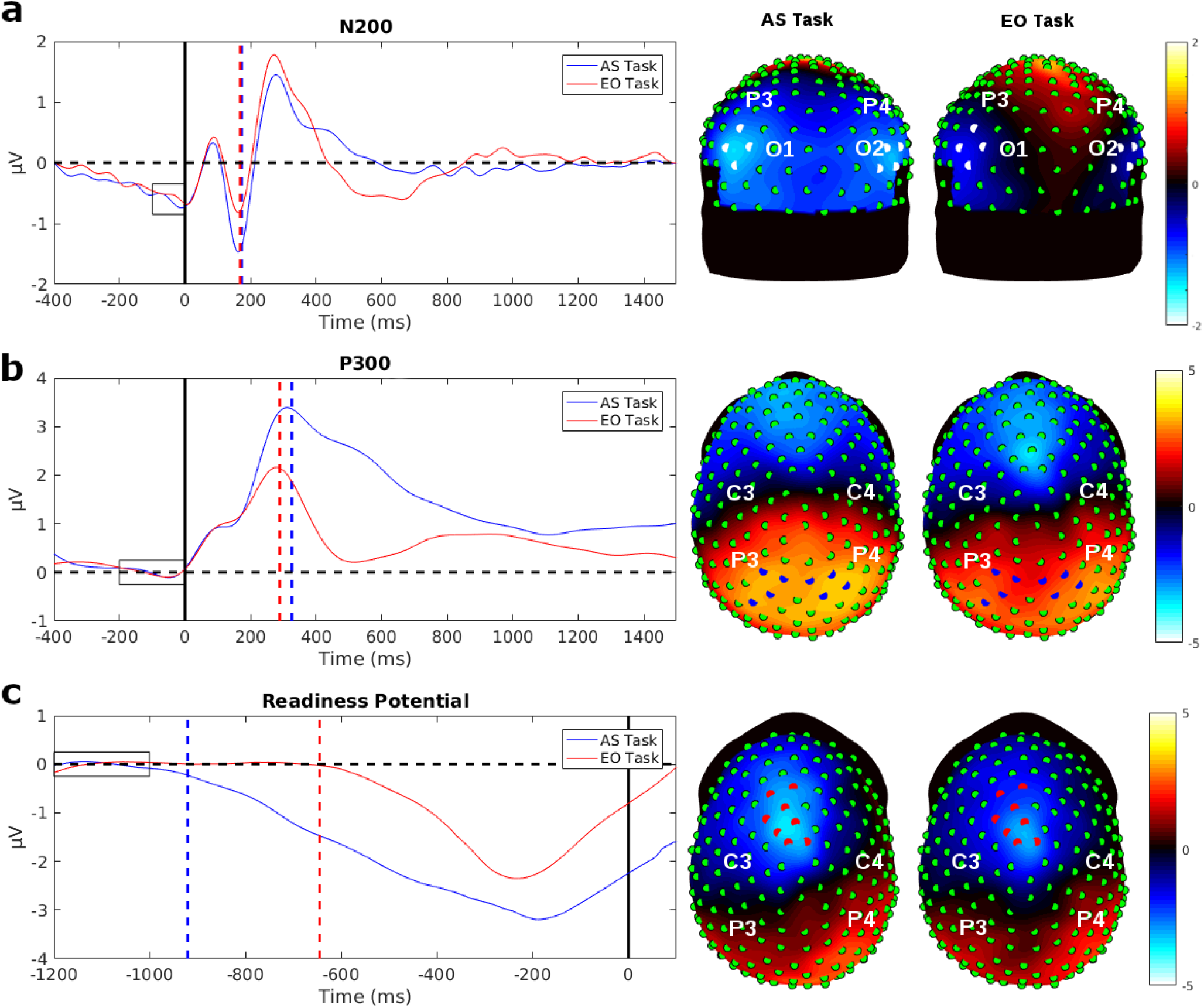
Evoked potential (EP) waveforms averaged across participants of the eight strongest electrodes for each EP in the AS task and the EO task. The dashed lines show the median peak latency or median duration time of the signals. The rectangles indicate the baseline interval. On each topographic map the location of these eight strongest electrodes are indicated and the map show the average potentials in a time window surrounding the peak. a) The N200 waveforms showed a bilateral distribution over occipital regions in both hemispheres. The N200 peak latency was nearly identical between the two tasks, with a median N200 peak latency of 173 ms in the AS task and 168.5 ms in the EO task. The topographies show mean potentials of the time interval from 100 ms to 150 ms, where the minimum peak amplitude was for both tasks. b) The P300 waveforms showed a bilateral distribution over parietal cortex for both tasks. The median P300 peak latency for the AS task was 326.5 and 289.5 ms in the EO task, respectively, showing about a 40 ms difference. The topographies show the mean potentials of the time interval from 240 ms to 340 ms, a window covering the peak latency in the two tasks. c) The response-locked RP waveforms showed strongest negativity over the midline area close to motor areas of the brain. The response-locked RP duration between the tasks varied greatly. The median response-locked RP duration in the AS task was 921 ms and in the EO task was 645 ms. In the AS task, the response-locked RP slowly ramped down to the negative minimum, compared to the EO task where the RP sharply ramped down to the minimum. The EEG topographies represent mean potentials of the time interval from −250 ms to −200 ms, where the minimum peak amplitudes occurred for both tasks.

### N200 and P300 peak latency estimate

The N200 and P300 showed greater individual difference in scalp topography as compared to the RP. In order to estimate the peak latency for each participant we first performed a singular value decomposition (SVD) on windowed filtered EP data (see Nunez et al., 2017 and Nunez et al., 2019a). For the N200, we used a window from 125 to 250 ms after stimulus onset, while for the P300, we used a window from 200 to 450 ms. For one participant with an extended P300, we used a window from 200 to 500 ms. In each participant’s data we verified visually that the largest SVD component had a scalp topography that matched the EP component and used the latency of the highest magnitude potential as the peak latency estimate.

### Statistical Tests

Analysis with linear models was performed on response-locked RP durations, stimulus-locked RP onset times, P300 peak latency, N200 peak latency, and RT distribution statistics. Mixed-effects ANOVA models were used to assess the effect of task with participants (and shoulder rotation for one model) as a random factor. In the RT tertile analyses, task and RT tertile were fixed factors, while participants were a random factor. Statistics reported are both F statistics and associated p-values as well as the Bayes Factors (BF), which describe the amount of evidence for a model with different means relative to a model with only one mean for both tasks or for RT tertiles (Rouder et al. 2012).

Paired samples t-tests were used to analyze differences in EPs between the two tasks, or between EPs in the same task (Student’s t-test for normally distributed data and Wilcoxon’s signed-ranked for non-normal data). Linear regression models were performed to assess the relationship between response-locked RP duration or P300 peak latency to RT or DT. In addition, a hierarchical linear regression model with participant treated as a random factor was also performed to analyze these same relationships. Conventional F-statistics on these models are presented. The amount of evidence for a model that has a non-zero regression slope over a model that has a regression slope of zero (Kass and Raftery 1995; Rouder et al. 2012) is presented as a BF. Adjusted *R*^*2*^ is also reported and describes the fraction of variance of the dependent variable (e.g. response time median) explained by the regressor variable. All other statistics were generated by either JASP, an open-source graphical software package for statistical analysis (JASP Team, 2017) or MATLAB (Natick, MA).

Bayes Factors (*BF1*) comparing models with regression slopes equal to one, indicating a one-to-one relationship, to models with unknown regression slopes were calculated by first fitting regression models (simple regressions and with participants as random factors) in JAGS (Plummer 2003) with wide priors on slope parameters (normal distribution centered on 1 with a standard deviation of 3) and then using the simple Savage-Dickey Density Ratio (Dickey and Lientz 1970; Wagenmakers et al. 2010). Note that Bayes Factors are sensitive to both models of comparison, and thus if the prior distribution (denominator) is compared to the posterior distribution (numerator), as we report with BF1, the BF will be highly sensitive to the prior probability even when the posterior itself does not change significantly (Liu and Aitkin 2008). For this reason, we caution over-interpreting BF1 which will increase with wider priors (smaller denominator) and decrease with less-wide priors (larger denominator). We also report parameter estimates given by the posterior medians and 95% credible intervals with the 2.5^th^ and 97.5^th^ percentiles of the hierarchical regression models that did not change significantly when using wider (normal distribution centered on 1 with a standard deviation of 1) and less wide (normal distribution centered on 1 with a standard deviation of 5) priors for the slope parameters.

### Surface Laplacian

The surface Laplacian was applied to the response-locked RPs to improve spatial resolution of the EEG. The surface Laplacian is the second spatial derivative of the EEG along the scalp surface and provides an estimate of the location of focal superficial cortical sources (Nunez and Srinivasan 2006). The scalp surface was modeled using the MNI-152 average head (Mazziota et al. 1995), and the surface Laplacian was calculated along a triangular mesh representing the scalp using a three-dimensional spline algorithm (see Deng et al. 2012).

## Results

### Behavioral Data: Response Time and Accuracy

RTs for the AS task (*Mean (M)* = 1031 ms, *Median (Md)* = 992 ms, *Standard Deviation (s)* = 139 ms) were much longer than the EO task (*M* = 627 ms, *Md* = 594 ms, *s* = 113 ms). A mixed-effects ANOVA model was used to analyze the RTs to estimate the effect of task with shoulder rotation and participants treated as a random factor. There was a significant effect of task on RTs (*F*(1,13) = 106.76, *p* = 0.02), with a decisive Bayes Factor (*BF* = 10^21^) supporting longer RTs in the AS task. There was no significant effect of direction of shoulder rotation; *F*(1,13) = 6.90, *p* = 0.15. Accuracy was very high for both the EO task (*M* = 0.99, *s* < 0.01) and the AS task (*M* = 0.97, *s* = 0.03).

### Drift-Diffusion Model: Decision and Non-Decision Time

We modeled the RT distributions with the DDM to separate NDT for perceptual preprocessing and motor execution from DT. The RT distribution for each participant is shown for each task in Fig. 2. The EO task was performed faster and with less variability across trials in each participant (Fig. 2, red histograms) compared to the AS task (Fig. 2, blue histograms). This result was expected since we did not expect the EO task to require much time for evidence accumulation and only necessitated perceptual preprocessing and motor execution. However, it may be true that some participants engaged in evidence accumulation on some trials in the EO task, as evidenced by the width and right tails of the distributions and non-zero DT estimates in the EO task. Posterior distributions of the NDT parameter were estimated for both tasks across participants (population level in the hierarchical model) and for each participant in each task. For each participant and task, a 95% credible interval of the NDT is presented by a shaded bar in Fig. 2. The fastest RTs in each task were good approximations of each participant’s NDT, confirming previous findings (see Ratcliff et al., 2016; Nunez et al., 2019a Fig. 5). The Pearson correlation between the 10^th^ RT percentiles and posterior medians of NDT for each participant was *ρ= 0*.*95* (*p* < *0*.*001*) in the AS task and *ρ= 0*.*93* (*p* < *0*.*001*) in the EO task. There was a wide variability in NDTs across participants but a relatively narrow 95% credible interval within each participant’s data.

**Fig. 4:**
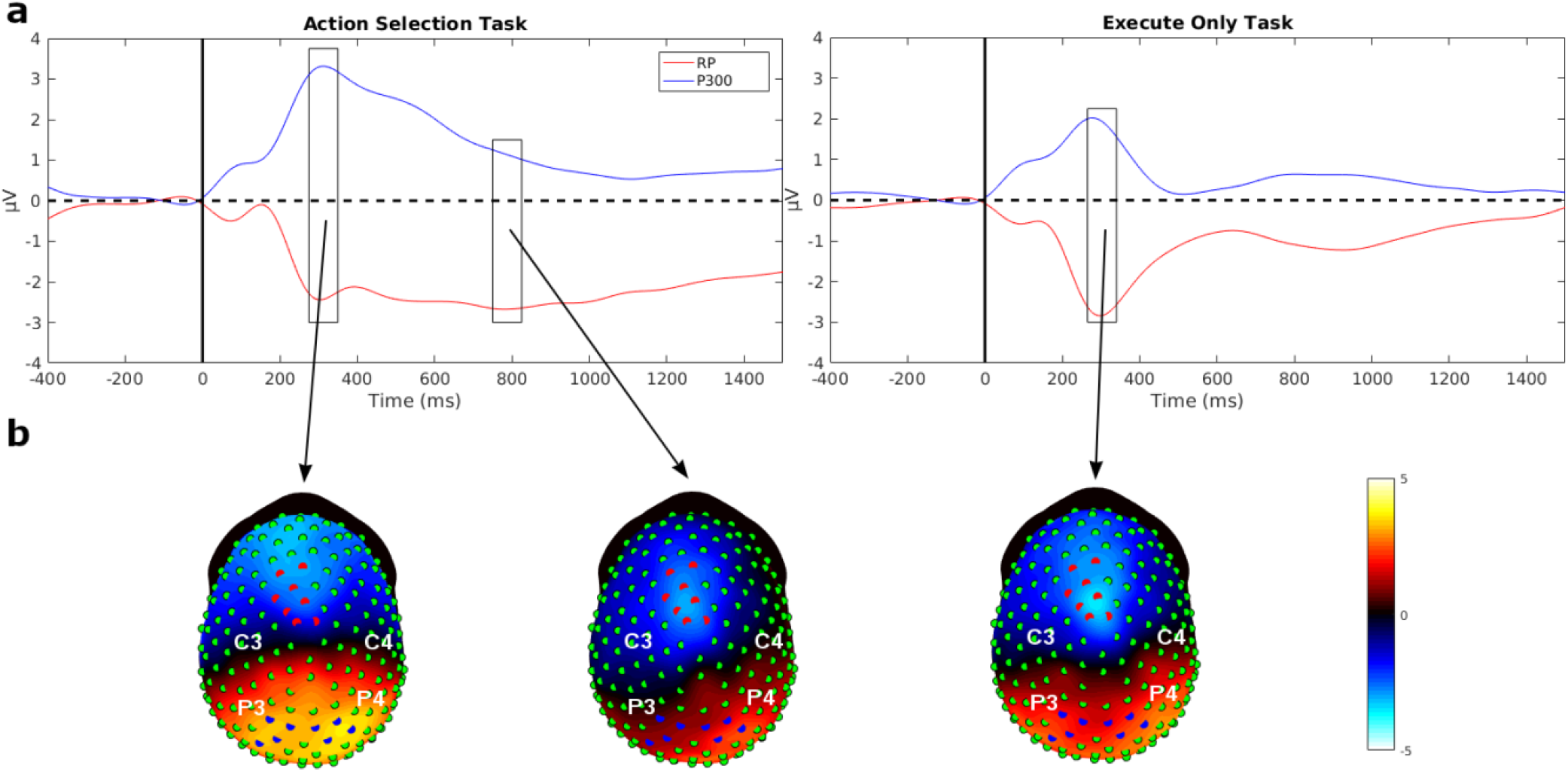
a) The time series of the stimulus-locked RP and the P300 averaged over the eight strongest channels. The stimulus-locked RP is in red and the P300 is in blue. The main difference seen between the two tasks are in stimulus-locked RP. In the AS task, after the stimulus-locked RP reached its minimum, there is a sustained negativity, whereas in the EO task, the signal has returned close to baseline. The P300 seemed to display similar time course in both tasks. b) In the AS task, the scalp topography is shown for the average potentials between 300 ms to 350 ms, revealing a topography similar to the one seen in Fig. 3b. When a later time period was averaged from 750 ms to 800 ms, the topographic distribution closely corresponded to the response-locked RP (see Fig. 3c). In the EO task, when averaging the potentials around the peak of the two signals, the topography looked similar to the response-locked RP (see Fig. 3c).

**Fig. 5:**
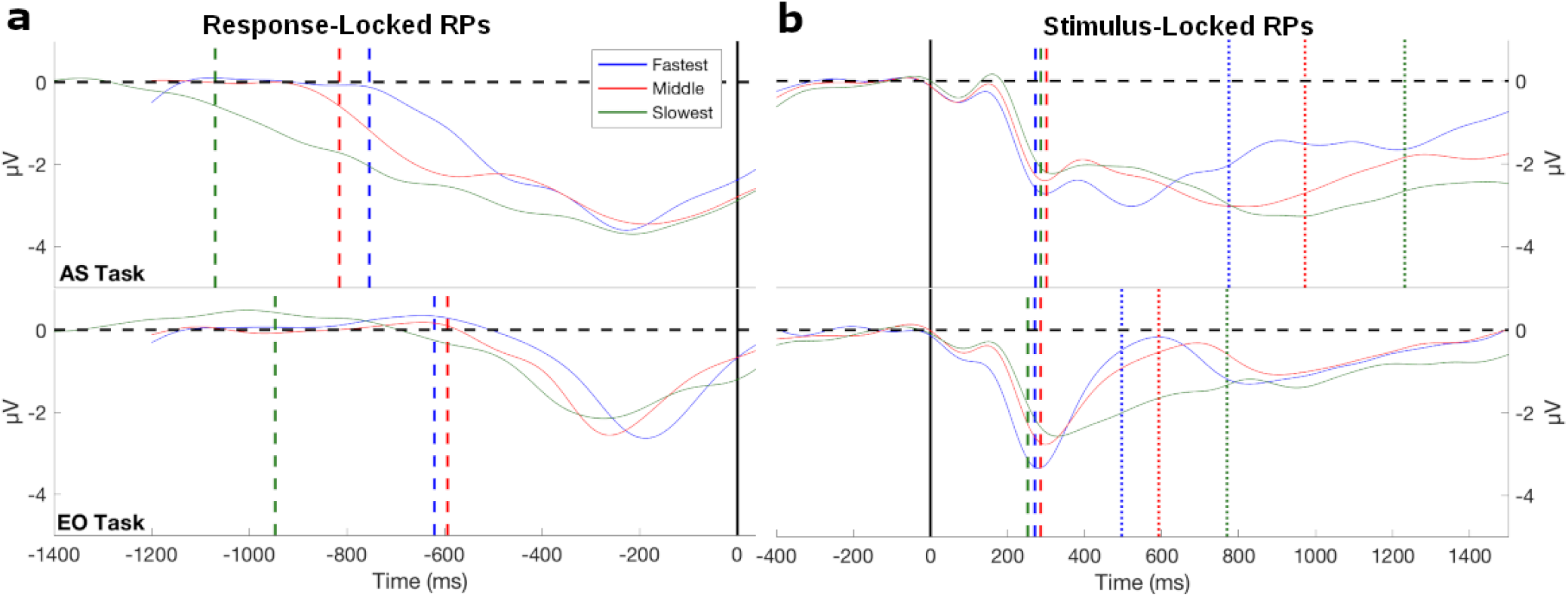
The RPs of the eight most negative motor channels averaged together and split into RT tertiles of fastest, middle, and slowest. a) The response-locked RPs for the fastest and middle tertiles are shown for a window of −1200 ms to 100 ms around the response indicated by a black vertical line. For the slowest RT tertile a longer window, −1500 ms to 100 ms, around the response, was used to accurately estimate response-locked RP duration. The dashed vertical lines represent the response-locked RP duration. In the AS task, the response-locked RP duration varied between conditions revealing the pattern of longer RTs having a longer RP duration. For the EO task, the fastest and middle condition had similar response-locked RP durations, and while the slowest was 300 ms longer. b) The stimulus-locked RPs are shown with a window of −400 ms to 1500 ms around stimulus onset indicated by the black vertical line. The stimulus-locked RP onset times were similar in both tasks for all tertiles as shown by the dashed vertical lines. The dotted vertical lines represent the average RTs in all of the conditions. One subject was excluded from this figure because of very fast RTs compared to the other participants. RP duration and onset times were estimated from this subject and included in the estimates of average duration and onset time and in the regression models shown in Fig. 7.

For the AS task, the median of the population-level posterior distribution of NDT was 377 ms with a 95% credible interval ranging from 317 ms to 438 ms. For the EO task, the median of the population-level posterior distribution of non-decision time was 288 ms with a 95% credible interval ranging from 235 ms to 338 ms. There was very small overlap (21 ms) found between the distributions for the two tasks indicating that the probability of the NDTs being the same between the AS task and the EO task was very small.

The DDM can be used to decompose RT into additive components of NDT and DT. For the AS task, the median RT was 992 ms, and the NDT estimate was 377 ms, indicating a typical DT of around 615 ms. In the EO task, the median RT was 594 ms, and the NDT estimate was 288 ms, indicating typical DT of approximately 306 ms. Thus, the longer RTs exhibited in the AS tasks (∼400 ms longer than EO) result from both longer DT (∼310 ms) and longer NDT (∼90 ms).

### Comparison of N200 and P300 peak latency between tasks

The N200 and P300 are stimulus-locked EPs recorded over occipital and parietal electrodes, respectively. Fig. 3 (a and b) shows the waveforms and topographic maps of each EP for each task averaged across all participants. The N200 shows the characteristic of negativity over occipital cortex at around 180 to 220 ms (Patel and Azzam 2005), and bilateral topography with the eight strongest electrodes in the average over participants distributed over both hemispheres. The N200 waveforms are averaged over the eight electrodes with strongest N200 indicated in the topographic map (Fig. 3a). The P300 is a positive potential recorded over parietal electrodes (Patel and Azzam 2005). The P300 waveform and topographic map is shown in Fig. 3b, with the waveform averaged over the eight strongest electrodes, distributed bilaterally over both hemispheres.

We examined the timing of the N200 in relation to the estimates of NDT estimates in each task obtained from the DDM. There was minimal difference in the N200 peak latencies between the two tasks at these eight electrodes (Fig. 3a). The AS task had a median N200 peak latency of 173 ms (*M* = 171.86 ms, *s* = 26 ms) and the EO task had a median N200 peak latency of 168.50 ms (*M* = 165.29 ms, *s* = 26 ms). N200 peak latencies were analyzed using a mixed-effects ANOVA, with participants treated as a random factor, indicating no significant difference found between the AS task and the EO task; *F*(1,13) = 3.72, *p* = 0.08, with *BF* = 1.16.

NDT consists of sensory encoding and response execution. In the behavioral data, we found the AS task has ∼90 ms longer NDT, while the N200 peak latency indicates no difference between tasks. Because N200 peak latencies has been shown to track sensory encoding time (Nunez et al., 2019a), this suggests that the difference between tasks in NDT is primarily due to response execution time. This might be expected as the AS task requires reprogramming a new response on each trial, while the EO task can be automated as the same response is required on each trial within a given block.

The median P300 latency in the AS task was 326.50 ms (*M* = 327.50 ms, *s* = 47 ms) and in the EO task was 289.50 ms (*M* = 294.64 ms, *s* = 44 ms). A mixed-effects ANOVA with participants treated as a random factor showed a significant difference between tasks; (*F*(1,13) = 12.58, *p* = 0.004, with *BF* = 10.47). The P300 peak latency was delayed by about 40 ms in the AS task as compared to the EO task, potentially reflecting the additional depth of processing required for stimulus categorization in the AS task versus stimulus detection in the EO task.

### Comparison of RP duration between tasks

We calculated the response-locked RPs for each participant to evaluate if the longer RTs in the AS task compared to EO task were reflected in the RP. The response-locked RP is a negative potential that begins prior to the response and reaches a negative peak at movement onset (Eimer 1998; Alexander et al. 2016). Fig. 3c shows the average across participants of the response-locked RPs, averaged across the eight channels with the strongest negative potential (selected from 256 channels) for both tasks. In our experiment, the button push corresponded to the completion of shoulder internal or external rotation, so the RP voltage minimum preceded the response marker by around 225 ms. This had little impact on the interpretation of the results, as the movement was identical in the two tasks and the response-locked RP reached a minimum at the same time around −250 to −200 ms in each participant and task; *t*(13) = 0.30, *p* = 0.77 with *BF* = 0.28. Fig. 3c shows the EEG topographies of the averaged potentials over the time period of the minimum peak (−250 ms to −200 ms). In these data, the response-locked RP showed a strong negative potential close to the midline and slightly left-lateralized.

The duration of the response-locked RP was defined as the time interval from the initial negative deflection of the response-locked RP (i.e., when the signal showed a consistent negative ramp), to the button press indicating completion of movement. The duration of the response-locked RP for the AS task (*M* = 870 ms, *Md* = 921, *s* = 88 ms) was longer than the duration of response-locked RP for the EO task (*M* = 680 ms, *Md* = 645, *s* = 146 ms) by approximately 190 ms. Response-locked RP duration was compared between tasks using a mixed-effects ANOVA with participants as a random factor. There was a significant difference in response-locked RP duration between the AS task and the EO task; *F*(1,13) = 18.72, *p* < 0.001, with *BF* = 304.24 indicating substantial evidence to support a longer duration of the response-locked RP in the AS task.

DT is defined as the difference between RT and NDT, and thus DT is an estimate of duration of the period of evidence accumulation in the DDM. The AS task has DT about 310 ms longer than the EO task. The P300 was delayed by about 40 ms which could account for only a small portion of the difference in DT. This difference may reflect the perceptual categorization in the AS task in comparison to stimulus detection in the EO task. In contrast, the duration of the RP was approximately 190 ms longer in the AS task than the EO task potentially indicating that response selection accounted for a larger portion of the increase in decision-making time.

### Comparison of Stimulus-locked RP to the P300

We examined the stimulus-locked RP at the same channels that showed the strongest negative potential for the response-locked RP for each participant in each task and compared it to the P300 recorded at parietal electrodes (Fig. 4). Similar to the response-locked RP, the stimulus-locked RPs showed a strong negative deflection at these electrodes. The onset of this negative potential occurred at a similar time for the AS task (*M* = 262 ms, *Md* = 246 ms, *s* = 76 ms) and the EO task (*M* = 270 ms, *Md* = 217 ms, *s* = 150 ms). A mixed-effects ANOVA for the stimulus-locked RP onset times of the negative deflection treating participants as a random factor showed no significant effect of task on onset times *F*(1,13) = 0.06, *p* = 0.81, with the Bayes Factor (*BF* = 0.36). This suggests highly ambiguous evidence regarding the difference between means. This also indicates that the onset of the RP precedes the P300 peak. Wilcoxon signed-rank test indicated that in the AS task, the stimulus-locked RP (*Md* = 246 ms) had an earlier onset time than the P300 peak latency (*Md* = 326.50 ms); *W* = 14, *p* = 0.01, with *BF* = 2.01. However, in the EO test, there was no significant difference between the stimulus-locked RP onset time (*Md* = 217 ms) and the P300 peak latency (*Md* = 289.50 ms); *W =* 28, *p* = 0.14, with *BF* = 0.69. This suggests that the response selection indexed by the RP and perceptual categorization indexed by the P300 take place in parallel in the AS task, consistent with previous studies showing covert EMG activity during decision-making (Servant et al. 2015), and studies showing the onset of the RP during decision-making (Ulrich et al. 1998; Gluth et al., 2013).

The striking difference between tasks was the extended duration of the negative potential in the AS task as compared to the EO task. Both tasks showed a negative potential that reaches a peak magnitude at roughly 300 ms after stimulus onset. Fig. 4b shows topographic maps of the potentials at selected time points. For the AS task, the stimulus-locked EPs at 300 ms post-stimulus is characterized by a strong positive potential over parietal channels, and a negative potential that is strongest at electrodes anterior to the eight channels with the strongest response-locked RP. This topography indicates that in this time window (about 300 ms after stimulus onset), the stimulus-locked RP incorporates the negative potential generated by the dipole sources of the P300. In contrast, by 800 ms after the stimulus, the topographic map indicated a strong negative potential with topographic distribution closely corresponding to the RP (see Fig. 3c) and very small positive potential over parietal channels, which have returned to nearly baseline (Fig. 4a). The EO task showed a topographic distribution at 300 ms, which incorporated both the negative potential at electrodes exhibiting the strongest RP and the positive potential over parietal cortex suggesting the two processes are entirely concurrent. The positive potential at parietal electrodes had a smaller magnitude and shorter duration.

We computed the average signal for the time period of 300 to 350 ms and 750 to 800 ms after stimulus onset for both the AS and EO task (see windows in Fig. 4a). A paired sample t-test was conducted to test the difference of the average of eight RP channels (Fig. 4a, in red) and the average of eight P300 channels (Fig. 4a, in blue) between the AS and EO task. In the interval 300 to 350 ms after stimulus onset, there was no significant difference found between the two tasks for the average potential of the RP channels; *t*(13) = 1.15, *p* = 0.27, with *BF* = 0.47. A significant effect was found between the two tasks when comparing the average of P300 channels in the same 300 to 350 ms interval; *t*(13) = 4.23, *p* < 0.01, with *BF* = 39.27. For the time period of 750 to 800 ms, the average of the eight RP channels was significantly different between tasks; *t*(13) = −3.44, *p* < 0.001, with *BF* = 11.25, while the average of the eight P300 channels was not significantly different in this time period; *t*(13) = 1.53, *p* = 0.15, with *BF* = 0.70.

### Stimulus-locked RP onset and Response-locked RP duration by Response Time Tertiles

We investigated if the RP onset (measured from stimulus-locked RP) or RP duration (measured from response-locked RP) was correlated to the RTs. For each participant, and in each task, the trials were sorted by RT and both stimulus and response-locked RPs were separately averaged for the trials in the slowest, middle, and fastest RT tertiles, as shown in Fig. 5.

For the AS task (Fig. 5a), the fastest RT tertile had the shortest RP duration (*M* = 754 ms, *Md* = 778 ms, *s* = 88 ms), then the middle RT tertile had a longer RP duration (*M* = 815 ms, *Md* = 851 ms, *s* = 109 ms), and the slowest RT tertile had the longest RP duration (*M* = 1070 ms, *Md* = 1098 ms, *s* = 142 ms). For the EO task, the fastest (*M* = 621 ms, *Md* = 635 ms, *s* = 146) and middle (*M* = 594 ms, *Md* = 526 ms, *s* = 182 ms) RT tertiles had similar RP duration while the slowest RT tertile (*M* = 947 ms, *Md* = 918, ms, *s* = 129 ms) had RP duration that was about 300 ms longer. A mixed-effects ANOVA was used to estimate the effect of task and RT tertile on RP duration treating participants as a random factor. There was no significant interaction between task and the tertile split (*F*(2,26) = 1.53, *p* = 0.24, *BF* = 1.60). There was a significant effect of task (*F*(1,26) = 23.08, *p* < 0.001) with evidence for a difference in mean duration times between tasks (BF = 71.81), and a significant effect of RT tertile (*F*(2,26) = 78.27, *p* < 0.001) with decisive evidence of a difference in mean duration between RT tertiles (*BF* =10^8^).

In contrast, the stimulus-locked RPs had onset times that were similar for the three RT tertiles in both tasks as shown in Fig. 5b. For the AS task, the fastest RT tertile had an average RP onset time at 273 ms (*Md* = 229 ms, *s* = 98 ms), the middle RT tertile had an average RP onset time of 301 ms (*Md* = 288 ms, *s* = 95 ms), and the slowest RT tertile had an average RP onset time at 287 ms (*Md* = 270 ms *s* = 94 ms). For the EO task, the fastest RT tertile had an average RP onset time of 271 ms (*Md* = 201 ms, *s* = 142 ms), the middle RT tertile had an average RP onset time of 286 ms (*Md* = 231 ms, *s* = 163 ms), and the slowest RT tertile had an average RP onset time of 253 ms (*Md* = 234 ms, *s* = 101 ms). The onset times of the RP did not account for differences in the response time tertiles. A mixed-effects ANOVA was used to estimate the effect of task and RT tertile on onset times treating participants as a random factor. There was no significant interaction between task and the tertile split (*F*(2,26) = 0.32, *p* = 0.73) with evidence for no interaction effect (*BF* = 0.03). There was no significant effect found in tasks (*F*(1,26) = 0.48, *p* = 0.50) with evidence for no effect of task (*BF* = 0.31), nor any effect by the RT tertiles (*F*(2,26) = 0.64, *p* = 0.54) with some evidence for no effect of RT tertile (*BF*= 0.17).

### P300 peak latency by Response Time Tertiles

The analysis by RT tertiles was repeated with P300 peak latency as shown in Fig. 6. When plotting the P300 signal by RT tertiles, the fastest RT tertile (*M* = 334 ms, *Md* = 333 ms, *s* = 45 ms), the middle RT tertile (*M* = 324 ms, *Md* = 325 ms, *s* = 45 ms), and the slowest RT tertile (*M* = 345 ms, *Md* = 326.50 ms, s = 61 ms) peaked at nearly the same time in the AS task (Fig. 6). In the EO task, the fastest RT tertile (*M* = 277 ms, *Md =* 286.50 ms, *s* = 33 ms) and the middle RT tertile (*M* = 292 ms, *Md =* 287 ms, *s* = 48 ms) had similar P300 median peak latency, while the slowest RT tertile (*M* = 332 ms, *Md* = 303 ms, *s* = 70 ms) showed a 40 to 50 ms longer latency compared to the other two conditions.

**Fig. 6:**
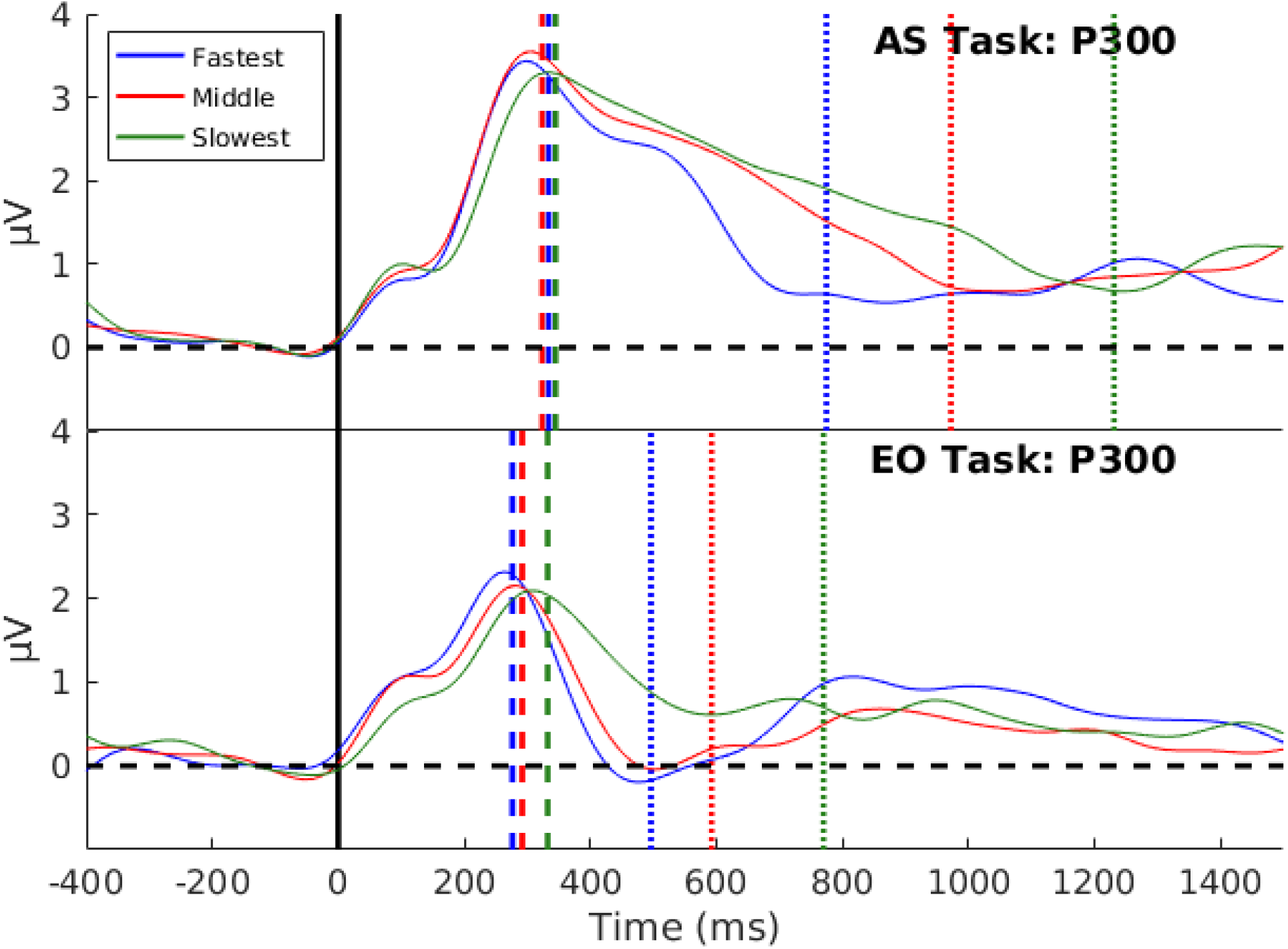
The P300 signal of the average of the eight most positive channels over parietal cortex are shown for the data split into RT tertiles in each task. In the AS task, the P300 peak latency is nearly the same in all of the conditions (shown by the vertical dashed line). In the EO task, it was similar in the fastest and middle condition, while the slowest condition showed a delay in peak latency of about 40 ms. This delayed peak in the slowest condition can possibly be due to a lack of attention and arousal during those trials due to the repetitive nature of the EO task. The average RTs for each condition in the two tasks are shown in the vertical dotted lines.

A mixed-effects ANOVA was used to estimate the effect of task and RT tertile on the P300 peak latency treating participants as a random factor. There was a significant interaction between task and RT tertile (*F*(2,26) = 6.30, *p* = 0.008, with *BF* = 4.48). Due to the interaction, a separate analysis was done for each task to examine the effect of RT tertile. In the AS task, there was no effect of RT tertile on P300 peak latency (*F*(2,26) = 1.76, *p* = 0.19, with *BF* = 0.55). However, a significant effect of RT tertile on P300 peak latency was found in the EO task (*F*(2,26) = 11.22, *p* < 0.001, with *BF* = 90.60). In the EO task, the slowest RT tertile showed a delayed P300. One explanation for this delayed peak in the slowest RT tertile is lack of attention and arousal during those trials due to the repetitive nature of the EO task. Another explanation could be the presence of an evidence accumulation process in the slowest trials and an absence of this process in the fastest trials.

### RP Duration as a Predictor of Response Time and Decision-Making Time

We tested if the duration of the response-locked RP was quantitatively related to RT and DT within each participant. A linear regression analysis was performed to determine if RP duration could predict RT for each task, using the RP duration and median RT computed for each RT tertile in each participant. For the AS task, RP duration was strongly correlated to median RT (*R*^*2*^= 0.50, *F*(2,40) = 40.43, *p* < 0.001), with decisive evidence of non-zero slope (*BF* = 10^4^). RP duration had a nearly one-to-one relationship with RT for this task (Fig. 8a) with *β* = .90 (*t*(40) = 6.36) and with some evidence of a regression slope of 1 (*BF1* = 15.45). The intercept of 217 ms reflects additional time required to account for RT, possibly reflecting visual stimulus processing time, and is consistent with the onset time of the negative deflection in the stimulus-locked RP (see Fig. 5b). In addition, a hierarchical linear regression model was fit to assess the same relationship by treating participants as a random factor that also assumed hierarchical mean parameters across participants. This model provided similar evidence with RP having a nearly one-to-one relationship with RT such that the hierarchical slope parameter was estimated near 1 (median posterior *β* = 0.99, 95% credible interval [0.78, 1.22], *BF1* = 25.31). The hierarchical intercept parameter in this model was estimated at 138 ms with some uncertainty (95% credible interval [-48 ms, 354 ms]).

**Fig. 7:**
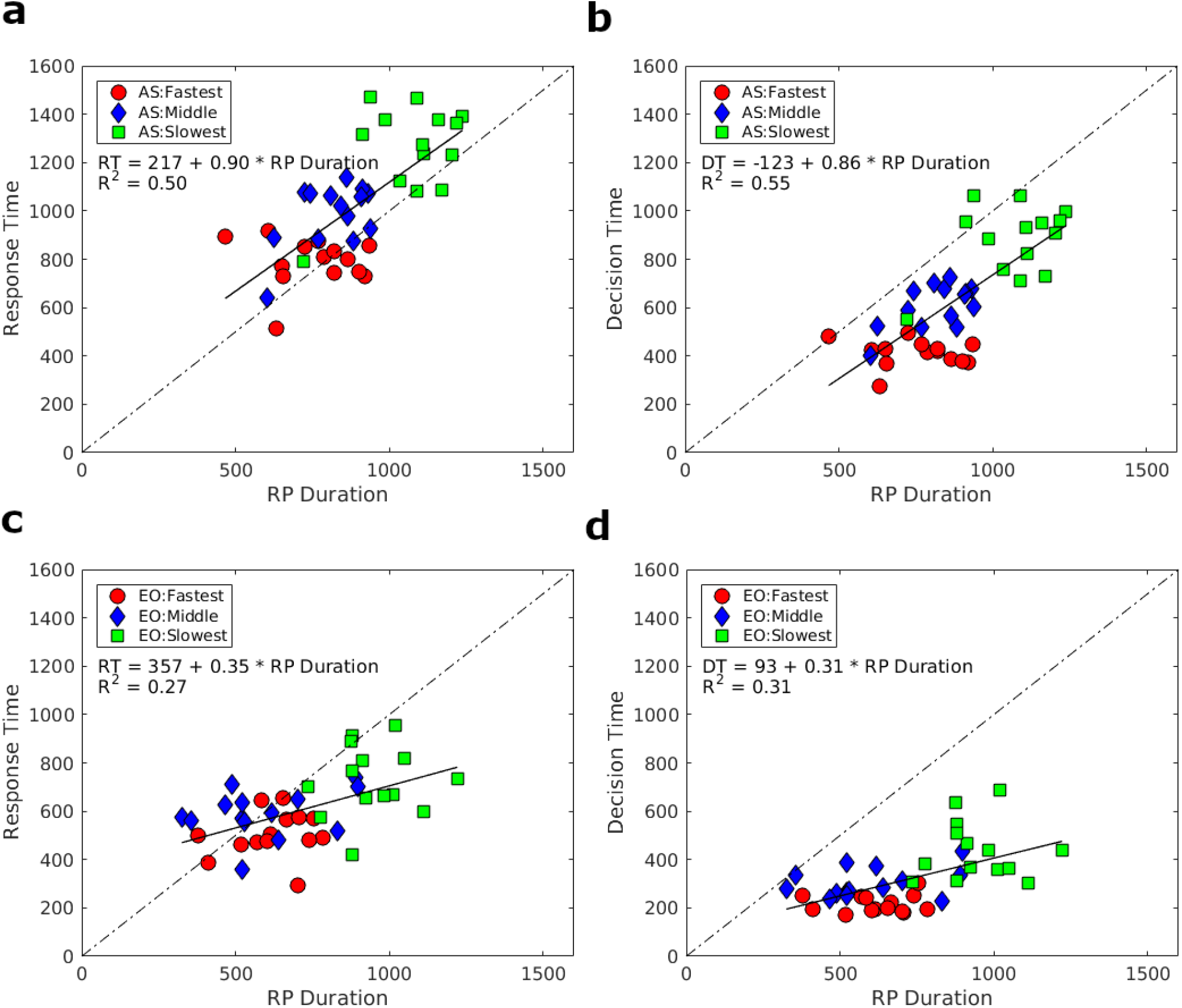
a) Regression model between response-locked RP duration and median RT for the AS task data divided into RT tertiles. b) Regression model between response-locked RP duration (absolute value of RP onset time) and median decision-making time for the AS task divided into DT tertiles. c) Regression model between response-locked RP duration (absolute value of RP onset time) and median response time for the EO task data divided into RT tertiles. d) Regression model between RP duration (absolute value of RP onset time) and decision-making time for the EO task divided into DT tertiles. This within participant effect showed that in the AS task, RP duration was strongly correlated to both RT and DT, with nearly a one-to-one relationship, indicating that the duration of RP was tracking decision-making process. For the EO task, the correlation was much weaker, and the slope not equal to one.

**Fig. 8:**
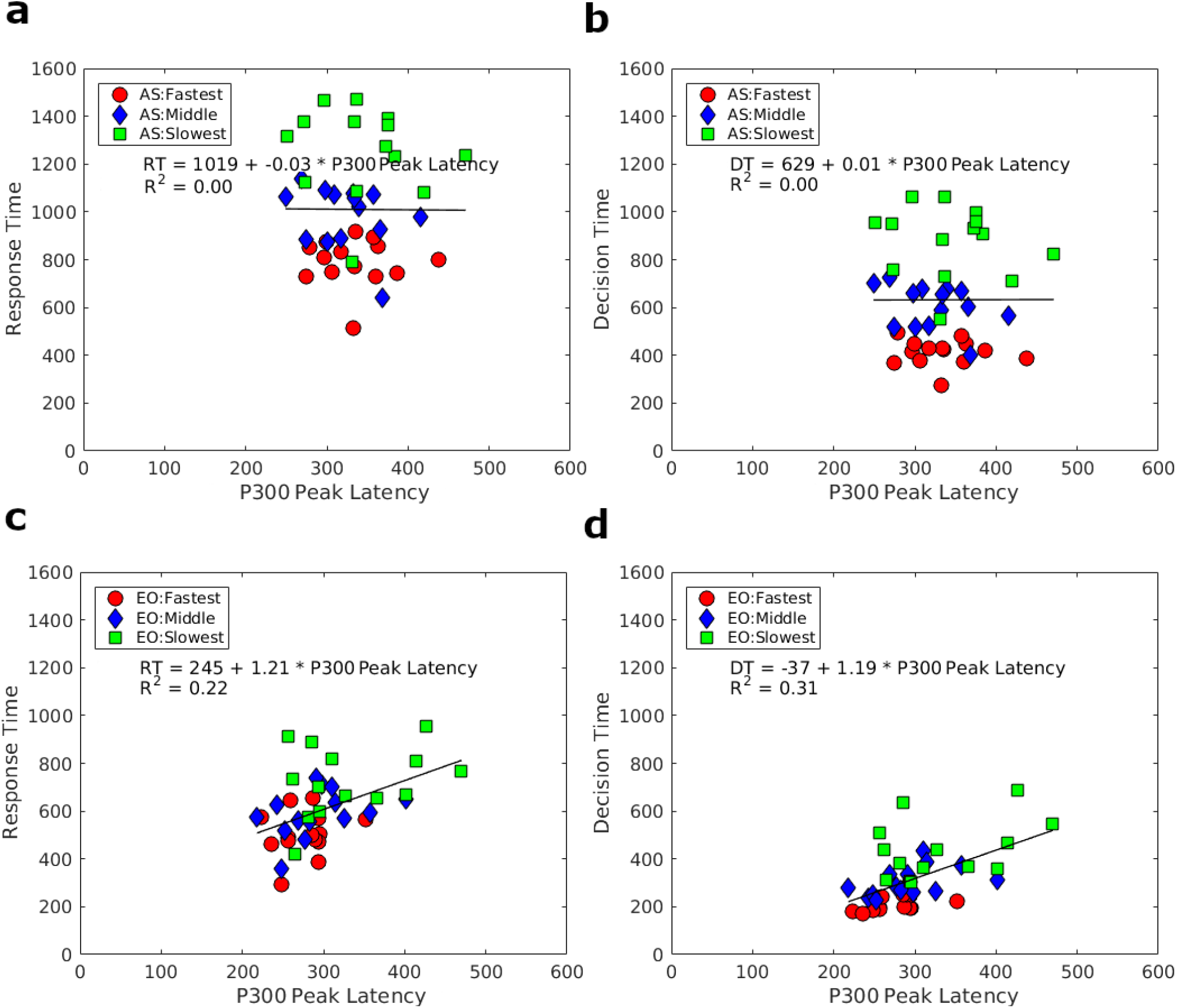
a) Regression model between P300 peak latency and median response time for the AS task data divided into RT tertiles. b) Regression model between P300 peak latency and median DT for the AS task divided into DT tertiles. c) Regression model between P300 peak latency and median response time for the EO task divided into RT tertiles. d) Regression model between P300 peak latency and DT for the EO task divided into DT tertiles. In the AS task, the P300 peak latency did not significantly correlate with either RT or DT. In the EO task, P300 peak latency was significantly correlated with RT and DT.

DT was calculated for each trial by subtracting the NDT (median of the posterior distribution of non-decision time estimated by the drift-diffusion model) from the RT on each trial. Similar to the RT tertile analysis, the trials were sorted by DT and EPs were estimated for tertiles of DT to estimate RP duration. The RP durations for DT tertiles were combined across participants to estimate a regression model.

In the AS task, the relationship between RP duration and median DT were fairly strong (*R*^*2*^ = 0.55, *F*(2,40) = 47.93, *p* < .001, *BF* = 10^5^), and the RP duration also had a near one-to-one relationship with DT (Fig. 7b) with *β* = 0.86 (*t*(40) = 6.92) and *BF1* = 4.04. The hierarchical linear regression model yielded similar results (median posterior *β* = 0.87, 95% credible interval [0.64, 1.11], *BF1* = 14.25). In contrast, for the EO task, the RP duration did not perform as well as a predictor of either RT or DT. There was a weaker (but significant) correlation between RP duration and RT (*R*^*2*^ = 0.27, *F*(2,40) = 14.60, *p* < .001), with *BF* = 60.16 (Fig. 7c) and a slope far less than one (*β* = 0.35, *t*(40) = 3.82 and *BF1* < 10^−59^). The hierarchical linear regression model yielded similar results (median posterior *β* = 0.36, 95% credible interval [0.21, 0.52], BF1 < 10^−211^). There was also a weak (but significant) relationship between RP duration and DT (*R*^*2*^ = 0.31, *F*(2,40) = 18.30, *p* < 0.001), with *BF* = 193.56 (Fig. 7d) and a slope far less than one (*β* = 0.31, *t*(40) = 4.28 and *BF1* < 10^−297^). The hierarchical linear regression model yielded similar results (median posterior *β* = 0.36, 95% credible interval [0.19, 0.46], BF1 < 10^−309^).

### P300 Peak Latency as a Predictor of Response-Time and Decision-Making Time

We tested whether P300 peak latency could predict RT or DT. In the AS task, the P300 peak latency had little to no correlation to the median RT tertiles (*R*^*2*^ = 0, *F*(2,40) = 0.001, *p* = 0.97, with *BF* = 0.30) as shown in Fig. 8a, and the hierarchical linear regression model of the same data yielded uncertain results with a wide posterior distribution for the slope parameter (median posterior *β* = 1.25, 95% credible interval [0.13, 2.33], *BF1* = 5.11). Also, no relationship was observed between P300 peak latency and DT tertiles (*R*^*2*^ = 0, *F*(2,40) < .001, *p* = 0.99), with *BF* = 0.30 as shown in Fig. 8b. Similarly, the slope of the hierarchical linear regression model of the same data was not significant as judged by the credible interval that overlapped zero (posterior median *β* = 0.57, [-0.33, 1.59], *BF1* = 4.03.

In the EO task, however, P300 peak latency did significantly correlate with RT (*R*^*2*^ = 0.22, *F*(2,40) = 11.25, *p* = 0.002), with *BF* = 19.73, with a slope a little more than one (*β* = 1.21, t(40) = 3.35 and *BF1* = 7.66) as shown in Fig. 8c. A hierarchical linear regression model of the same data estimated a hierarchical slope with a posterior median of *β* = 1.42 (95% credible interval [0.67, 2.12], *BF1* = 4.79). There was a stronger relationship with DT and P300 peak amplitude that also showed a significant correlation (*R*^*2*^ = 0.31, *F*(2,40) = 17.80, *p* < 0.001, with *BF* = 165.02. There was a slope of slightly >1 (*β* = 1.19, *t*(40) = 4.22 and *BF1* = 11.05) as shown in Fig. 8d. A hierarchical linear regression model of the same data estimated a hierarchical slope with a posterior median of *β* = 1.20 (95% credible interval [0.65, 1.80], *BF1* = 8.23). As noted earlier, in the EO task, the slowest tertile of RT was associated with a delayed P300, which could possibly be an effect of arousal rather than decision-making or a reflection of some evidence accumulation in the slowest trials.

### Surface Laplacian analysis of the RP

The RPs in each participant were processed with a surface Laplacian to identify superficial focal current sources. In both tasks, with the application of a surface Laplacian, the current density estimates were localized over the midline somewhat anterior to bilateral motor cortex. Three electrodes were positioned over the strongest signals, and the time course shows that of those three, one electrode over the right midline area showed a positive signal while two electrodes over the left midline area showed a negative signal in both tasks (Fig. 9a). This suggests that the RP involves a lateralization of current density in areas of the motor system that generate the RP and is discussed further in the Discussion. Fig. 9b shows the topographies of the surface Laplacian averaged over −250 ms to −200 ms before the response, and the three electrodes with the highest magnitude current density are marked in green, red, and blue and labeled 1-3, corresponding to the waveforms in Fig. 9a. The overall pattern shows that the right hemisphere exhibited more positive current density in comparison to the left hemisphere, for both of these tasks involving right arm movement.

**Fig. 9:**
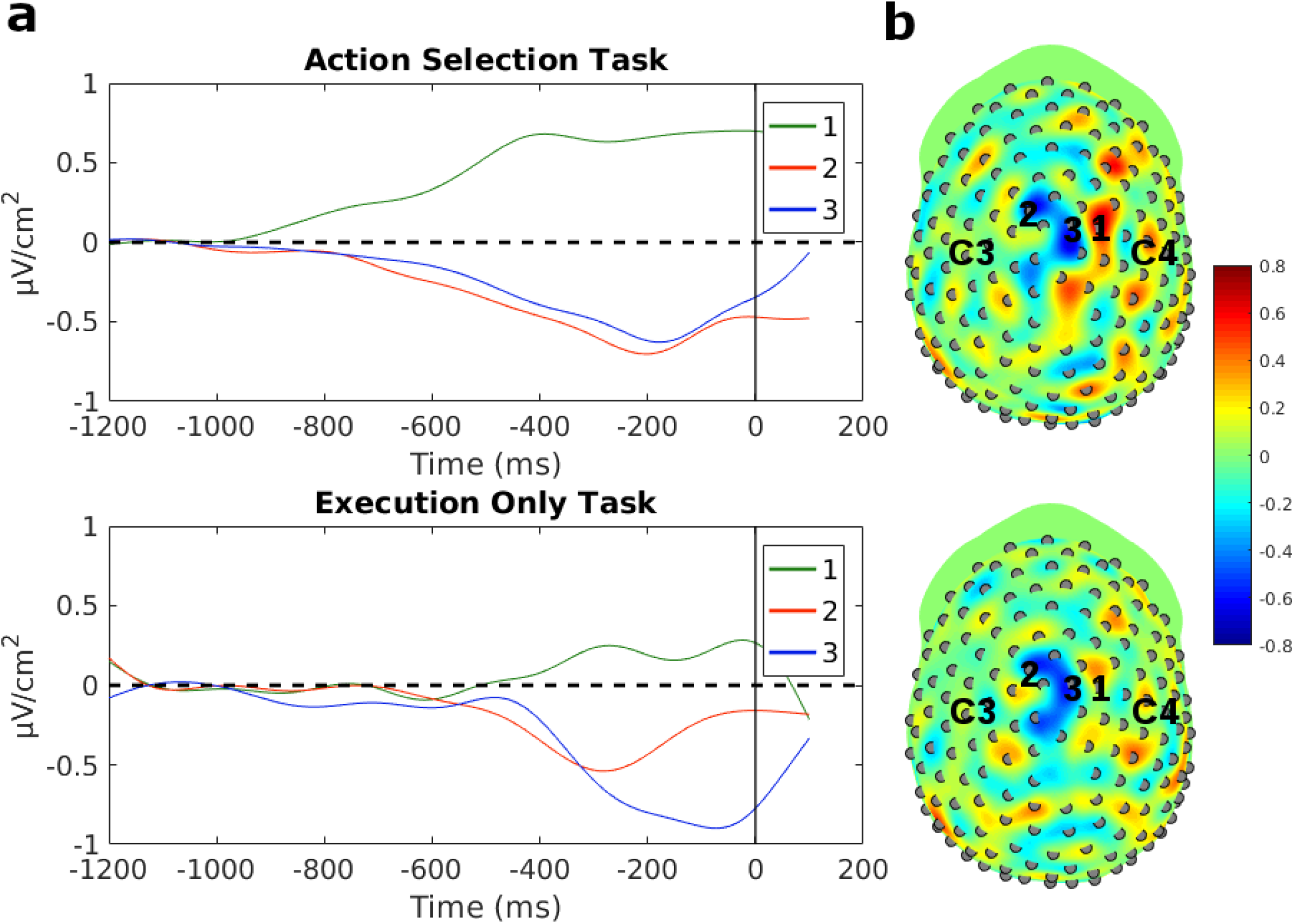
By applying a surface Laplacian, the strongest current density was localized close to the midline, potentially generated by bilateral structures in the motor cortex close to the midline. There were two electrodes that showed strong negativity over the left midline area and one electrode that showed strong positivity over the right midline area that could be suggestive of lateralization of the motor system. a) The time course of the three electrodes that generated the strongest current source density. b) The topography of current source density with the surface Laplacian applied. The three midline electrodes with greatest activity are marked in red, green, and blue and labeled 1-3. For reference, C3 and C4 electrodes are labeled.

## Discussion

Evidence accumulation is believed to be the underlying mechanism of decision-making, determining the time it takes to select between alternative decisions (Link and Heath 1975; Ratcliff and McKoon 2008). Previous studies have shown that parietal cortex activity exhibits characteristics of evidence accumulation (Shadlen and Kiani 2013; Kelly and O’Connell, 2013; O’Connell et al. 2018). These studies have modeled the mechanisms of perceptual categorization in decision-making, followed simply by the initiation of response execution. In this study, we made use of the AS task (O’Shea et al. 2007) to investigate the contribution of response selection to decision-making. RT and accuracy data were fit to a DDM (Ratcliff and McKoon 2008) in order to partition RT into NDT (for stimulus encoding and motor execution) and DT (corresponding to the duration of evidence accumulation). We found that, for the AS task only, the duration of RP recorded over motor-related areas of the brain has a one-to-one relationship with DT. The results indicate that in the AS task, *response selection* can be modeled with an evidence accumulation process reflected in the timing of the RP.

### Readiness Potential reflects decision-making in the Action Selection task

RPs display the characteristic of slow ramping negativity reaching a minimum at the start of movement, resolving to baseline as the movement completed and has been characterized to reflect response selection and multiple movement-related processes (Rohrbaugh and Gaillard 1983; Osman et al. 1995; Eimer 1998; Miller et al. 1999; Leuthold et al. 2004; van Boxtel and Böcker 2004; Alexander et al. 2016). The duration of the RPs (absolute value of the RP onset time) from the AS task was on average about 190 ms longer than the duration of the RP from the EO task. The stimulus-locked RPs at the same channels exhibited a negative ramp that onset identically at around 265 ms for both tasks, suggesting that the onset of the RP takes place after visual encoding but before evidence accumulation. Moreover, the onset of the RP precedes the P300/CPP peak, which has been suggested to be an indicator of evidence accumulation reaching a boundary (O’Connell et al. 2012). Past work did not find evidence for an effect of stimulus intensity on RP duration (Miller et al. 1999), indicating that the RP is independent of perceptual processing. Although other studies have found that stimulus intensity does influence motor execution time (Resulaj et al. 2009; Buc Calderon et al. 2015; Dotan et al. 2018; Weindel et al. 2020), here we found that the RP does account for timing of decision-making in a response selection task, although the RP likely also reflects motor preparation and execution.

To clarify if RP quantitatively tracked DT, a linear regression analysis was performed with the tertile split of trials by RTs and DTs. Showing a within participant effect, in the AS task, RP duration and RT had a strong relationship, indicating a nearly one-to-one relationship (*BF1* = 25.31 from hierarchical regression model) between increasing duration of RP and RT (Fig. 6a). The intercept of the standard regression model was 217 ms, which was consistent with time for perceptual processing (Thorpe et al. 1996; Nunez et al. 2019a), and with the onset of the stimulus-locked RP at about 265 ms (note that the 265 ms estimate is the center of an interval from 203-328 ms where the derivative of the RP waveform is consistently negative). RP duration and DT also had a linear relationship (Fig. 6b) indicating a close one-to-one relationship (*BF1* = 14.25 from hierarchical regression model) between RP duration and DT. These models provide strong evidence that RP duration tracks DT, and that the intercepts of the regression model with RT accounted for the visual encoding time prior to the onset of evidence accumulation consistent with the onset time of the stimulus-locked RP.

In the EO task, RP duration did not predict DT and RT as well as it did in the AS task. A lack of variability in the EO task RTs, compared to AS task RTs, may partially explain this finding, as shown in Fig. 2. Moreover, in the EO task, the decision of which action to execute is fixed. As a consequence, variability in RT or DT may be more strongly influenced by variability due to stimulus processing (detection) or response execution rather than in decision-making.

### The P300 does not track decision-making in the Action Selection task

In previous studies, the central-parietal positivity (CPP) and P300 amplitude and timing of peak positive potential have been found to be reflective of evidence accumulation process towards a decision (O’Connell et al. 2012; Kelly and O’Connell 2013; Twomey et al. 2015). In vigilance tasks (specifically gradual reduction in contrast and motion detection task with different coherence levels), the peak of CPP was delayed as RT increased (O’Connell et al. 2012; Kelly and O’Connell 2013). As the difficulty of identifying and processing the stimulus increased, there was a delay in the CPP, with a peak right before motor execution. This was attributed to longer decision-making since the peak of the CPP occurred at motor execution. However, some of this delay may also reflect increased time for perceptual processing, as these studies employed stimuli at varying stimulus-to-noise ratios. The task we have used in this study employed simple shapes presented without any noise, and the decision-making involves a significant component of response selection rather than perceptual categorization. The P300 was identified in each task, but the time of the peak was delayed by about 40 ms in the AS task, despite a nearly 400 ms difference in RTs (Fig. 4c) and a 310 ms difference in DTs. Moreover, when the data were split into tertiles of fastest, middle, and slowest RTs, the P300 peak latency did not predict RTs or DTs in the AS task within participants. We also did not find a separate CPP from the P300 that predict RT or DT that better matched the CPP as found in other studies (O’Connell et al. 2012; Kelly and O’Connell 2013; Rangelov and Mattingley 2020). This could be due to different task structure and demands.

### Evidence accumulators in the human brain

Decision-making includes both perceptual categorization and response selection. Our results suggest that by monitoring both perceptual and motor signals, EEG data can be used to understand the distinct contributions of perceptual and motor decision-making to RT data depending upon the task demands. The CPP or P300 may reflect an evidence accumulation process for perceptual categorization, which have been found to be closely related to evidence accumulation in tasks with challenging perceptual decision-making (Philiastides et al. 2006; Ratcliff et al. 2009; O’Connell et al. 2012; Kelly and O’Connell 2013). In the AS task, perceptual categorization was not decisive in determining RT and the decision-making has a large component of response selection. Our findings indicate that the RP reflects evidence accumulation in the motor system for response selection, which is decisive in determining RT in this task.

RT and choice behavior during visual decision-making tasks are well characterized by models that assume a continuous stochastic accumulation of evidence (Link and Heath 1975; Usher and McClelland 2001; Ratcliff and McKoon 2008; Brown and Heathcote 2008; Ratcliff et al. 2016). Studies in animal models showing increasing firing rates in parietal cortex during perceptual decision-making (Roitman and Shadlen 2002; Huk and Shadlen 2005, Churchland et al. 2008), motivated the earlier studies of the P300 in parietal cortex (Philiastides et al. 2006; Ratcliff et al. 2009; O’Connell et al. 2012; Kelly and O’Connell 2013) and the present study of the RP over motor-related areas of the brain. However, the relationship between progressively increasing firing rate and ramps in slow-wave EEG potentials are complicated by a number of factors. EEG potentials reflect synchronous synaptic potentials at a macroscopic (cm) scale (Nunez and Srinivasan 2006). The sources of the EEG are the currents in extracellular space − excitatory postsynaptic potentials (EPSPs) generate positive sources inside the membrane and negative current in the extracellular space while inhibitory postsynaptic potentials (IPSPs) generate negative sources inside the membrane and positive current in extracellular space. Thus, increasing firing rates might be expected to generate increased negative potentials as negative extracellular potentials reflect (locally) more EPSPs, suggesting that a negative ramp is related to increased cortical excitability facilitating firing of action potentials. However, we are cautious about this interpretation of the RP because of the physics of EEG recording. Scalp potentials are recorded at a distance from the cortex, and thus dominated by dipole components of the brain current source distribution (Nunez and Srinivasan 2006; Nunez et al. 2019b). Thus, the sign of the observed potential may merely reflect whether the negative or positive pole of the dipole is closer to the scalp. Despite this complication, it is reasonable to interpret a slow ramp in EEG potentials (of either sign) as a correlate of progressive change in firing rates, providing a means to investigate accumulator processes in the human brain.

The RP duration in the EO task was somewhat surprising because if it were an evidence accumulator, we would not expect it to be longer than RTs for this task. Furthermore, the only case where waveforms begin before stimulus onset is the RP estimate for the EO task. There are several possible reasons for observing these effects. One is the automatic nature of the task. In the study by Libet and colleagues (Libet et al. 1983) it has been suggested that response selection, indexed by the RP, precedes the conscious decision to move. In the EO task, despite the introduction of random onset times, there is essentially a repetitive motion that the subject may prepare for prior to stimulus presentation. Another is that the RP has been observed before stimulus onset in cases where there is pre-cued information given about the response to be made (Osman et al. 1995; Leuthold et al. 1996; Ulrich et al. 1998; Leuthold et al. 2004). This could explain why we only see this in the EO task because participants already know which direction they will be responding.

### Sources of the Readiness Potential

The scalp topographies of the RP reveal that the strongest negative potentials occur over motor areas close to the midline and anterior to C3/C4 (Fig. 3b). The Bereitschaftspotential for finger movements is composed of an early shallow ramp over midline areas, such as supplementary motor areas (SMA) and pre-SMA, and a steeper negative slope over contralateral motor areas (corresponding to C3/C4) that reaches a negative minimum at movement onset (Libet et al. 1983; Shibasaki and Hallett 2006). In our task, which involved shoulder internal and external rotation movements, the RP had a focus entirely over midline areas from RP onset to movement onset, consistent with activity in the shoulder representation in primary motor cortex which is located close to the midline (Penfield and Rasmussen 1950). The higher spatial resolution of the surface Laplacian (Fig. 9) clearly indicated a focal source close to the midline, with opposite polarities between hemispheres. The lateralization found in the surface Laplacian, with positive potential in the hemisphere ipsilateral to movement and negative potential in the contralateral hemisphere, has been thought to reflect response inhibition and activation patterns and has also been observed in other choice reaction time tasks (Burle et al. 2004).

#### Limitations of the study

This study has a few methodological limitations that need to be acknowledged. There was no counterbalancing in the order of administration of the two tasks, with all the participants starting with the AS task, thus ending with the EO task. This might have contributed to some fatigue effect in the last EO block for that specific shoulder rotation, influencing the measures of their RTs. With the exploratory nature of the task, a formal power analysis was not conducted to determine the optimal sample size; future confirmatory studies should perform such an analysis to ensure sufficient sample size. The relationship of the RP to DT should also be explored in tasks with more difficult response selection. In both the EO and AS task, participants were very accurate (all participants were 100% accurate in the EO task and were between 88% and 100% accurate in the AS task with a mean of 98% accuracy).

## Conclusion

Decision-making has several underlying components including stimulus encoding, perceptual categorization, response selection, and response execution. In this study, we made use of a task where decision-making mostly involved response selection. For this task, we find that the duration of a signal linked to the motor system, the RP, has a one-to-one relationship with amount of time required to make the decision, which is modeled by a stochastic evidence accumulation process. This close relationship between the RP and the evidence accumulation process supports the notion that an accumulator process is a general neural implementation of decision-making in both sensory and motor systems, both when individuals take actions in response to stimuli (Shadlen and Kiani 2013) or of their own free will (Schurger et al. 2012). Our results suggest that the contributions of sensory or motor systems to variability in decision-making time can be assessed by the timing of sensory and motor evoked potentials.

## Acknowledgements

This research was supported by grants from the NSF to JV and RS (1658303 and 1850849), grants from the NIH to JMC (K99HD091375 and T32AR047752) and SC (K24HD074722), and a grant from the University of California, Irvine Institute for Clinical and Translational Science (UL1-TR000153). Kiana Scambray is thanked for her help with data analysis in studies related to this paper. Jennifer Wu and Vu Le are thanked for their help on task design and construction of splint apparatus to record responses for this paper.

## Data and code availability

Artifact-correct EEG data is available upon request to the corresponding author: MATLAB and JAGS analysis code are available on https://osf.io/7r6af/ and in the following repository https://github.com/mdnunez/RPDecision (as of July 2020).

## Notes

### Competing Interest Statement

The authors have declared no competing interest.

