## Supplemental Tables for "Timing of readiness potentials reflect a decision-making process in the human brain"

---

---

### 1. Supplementary Tables

Variance of correct-RT percentiles explained by in-sample prediction

| | **Action Selection (AS)** | **Execution Only (EO)** |
|---|---|---|
| **10th RT Percentile** | 95 % | 98 % |
| **30th RT Percentile** | 95 % | 97 % |
| **50th RT Percentile** | 98 % | 95 % |
| **70th RT Percentile** | 98 % | 91 % |
| **90th RT Percentile** | 96 % | 79 % |

Table 1: Percentage of across-participant variance explained by in-sample prediction ($R^2_{\text{pred}}$, see Equation 2 in Nunez et al., 2017) by the hierarchical DDM for accuracy and percentiles of correct-RT distributions. Note that participants made very little errors in either condition, and therefore accurately evaluating the model's descriptive ability of error-RT percentiles was not possible. Because the hierarchical DDM describes fast RT percentiles well in both conditions, we trust the non-decision time (NDT, $\tau$, see Nunez et al., 2019) and decision-time (DT) estimates estimated by the median posteriors of $\tau$ parameters in each condition.

Hierarchical DDM parameter estimates

| | **AS Mean (95% CI)** | **AS Part. Range** | **EO Mean (95% CI)** | **EO Part. Range** |
|---|---|---|---|---|
| $\tau$ **(sec)** | 0.38 [0.32, 0.44] | 0.24 to 0.49 | 0.29 [0.23, 0.34] | 0.11 to 0.45 |
| $\alpha$ **(evid.)** | 3.92 [3.39, 4.44] | 3.85 to 4.11 | 2.96 [2.55, 3.41] | 2.88 to 3.16 |
| $\delta$ **(evid./sec)** | 3.17 [2.69, 3.64] | 2.56 to 4.36 | 4.76 [4.24, 5.32] | 3.15 to 6.17 |
| $\lambda$ **(probability)** | 5.32% ** | 1.01% to 20.49% | 2.30% ** | 0.78% to 5.16% |

Table 2: Estimates for hierarchical mean parameters $\mu$ for the Action Selection (AS) and Execution Only (EO) conditions are given by the posterior medians (1st and 3rd columns). 95% Credible Intervals (CI) are also shown and were calculated using the 2.5th and 97.5th percentiles of the posterior distributions (1st and 3rd columns in brackets). Also shown are the participant (Part.) ranges for each parameter given by the minimum and maximum posterior medians of participant-level parameters (2nd and 4th columns). Non-decision time $\tau$ estimates are in seconds (sec). Boundary separation $\alpha$ estimates are in evidence units (evid.). Drift rate $\delta$ estimates are in evidence units per second (evid./sec). Lapse parameters are given by probabilities. **Note that the lapse probability mean is the mean of the posterior medians of participant-level parameters because our model did not include hierarchical lapse parameters.
